# Primary and secondary motoneurons use different calcium channel types to control escape and swimming behaviors in zebrafish

**DOI:** 10.1101/2020.07.14.202853

**Authors:** Hua Wen, Kazumi Eckenstein, Vivien Weihrauch, Christian Stigloher, Paul Brehm

**Author notes:** Corresponding Author: Paul Brehm, Oregon Health & Science University, 3181 S.W. Sam Jackson Park Rd., Portland, OR 97239. University of California at Irvine, Irvine, CA 92697.

## Abstract

The escape response and rhythmic swimming in zebrafish are distinct behaviors mediated by two functionally distinct motoneuron (Mn) types. The primary (1°Mn) type depresses, has a large quantal content (Qc), and a high release probability (Pr). Conversely, the secondary (2°Mn) type facilitates and has low and variable Qc and Pr. This functional duality matches well the distinct associated behaviors, with the 1°Mn providing the strong, singular C-bend initiating escape and the 2°Mn confers weaker, rhythmic contractions. Contributing to these functional distinctions is our identification of P/Q type calcium channels mediating transmitter release in 1°Mns and N type channels in 2°Mns. Remarkably, despite these functional and behavioral distinctions, all ~15 individual synapses on each muscle cell are shared by a 1°Mn bouton and at least one 2°Mn bouton. This novel blueprint of synaptic sharing provides an efficient way of controlling two different behaviors at the level of a single postsynaptic cell.

## Introduction

The complex body plan associated with most classes of vertebrate animals provides for a wide repertoire of movements. This involves a large number of different specialized muscle groups that are separately innervated by different spinal Mns. By contrast, fish largely conform to a simplified organization reflected in a segmentally repeating pattern of axial muscle. Despite the simplicity and limitations of this organization, larval zebrafish are still capable of mounting distinct swimming behaviors that, through necessity, have evolved to use the same fast skeletal muscle cells. The principal behaviors correspond to a C-bend, which mediates escape, and rhythmic swimming (Fetcho and Faber, 1988; Muller and van Leeuwen, 2004). The C-bend is likened to an all-or-none behavior, reflecting a single powerful stereotypic contraction. The ensuing rhythmic swimming covers a range of speeds that reflect differential power development among the same peripheral muscle cells. The C-bend is the most-well studied and best understood, due to both its simplicity and predictability (Fetcho and Faber, 1988; Korn and Faber, 2005). The rhythmic swimming, however, involves control mechanisms that appear to govern contraction strength within and among muscle cells; a process that is considerably more complex.

It is generally agreed that the C-bend is mediated by firing of the four large 1°Mns located in each spinal cord hemi-segment (Liu and Westerfield, 1988; Fetcho and O’Malley, 1995). By contrast, the rhythmic swimming reflects the distributed firing among the ~40 2°Mn types within each hemi-segment (Myers et al., 1986; Westerfield et al., 1986). The principal control over rhythmic swimming falls to the 2°Mns with 1°Mns participating at only the fastest swim speed (Liu and Westerfield, 1988; McLean et al., 2007). Unlike the stereotypic C-bend, fine-tuning of swim speed involves a size-dependent recruitment among the 2°Mns (McLean et al., 2007; Gabriel et al., 2011). Firing among the smallest, most ventral 2°Mns is causal to slow swimming whereas recruitment of more dorsal, larger size 2°Mns results in faster swimming (McLean et al., 2007). For the most part, studies on the mechanisms underlying Mn size dependence of swim speed have been focused on distinctions in premotor spinal circuitry and intrinsic firing properties among the Mns (McLean et al., 2007; Gabriel et al., 2011; Menelaou and McLean, 2012). However, there is growing appreciation of peripheral mechanisms that also contribute to the regulation of power development. The first mechanism relates soma size to overall peripheral branching. A more extensive branching on the part of larger 2°Mns likely reflects a greater number of muscle cells receiving innervation (Bello-Rojas et al., 2019), a proposition further supported by the present study. Accordingly, recruitment of larger Mns would result in increased power by activating a greater number of muscle cells. A second, and novel peripheral determinant of swim speed, was revealed through paired patch clamp recordings of 2°Mns and target muscle (Wang and Brehm, 2017). The recordings revealed a direct relationship between quantal content (Qc) and 2°Mn size at the level of individual muscle cells. Consequently, a larger Qc would aid in contraction strength by producing a greater depolarization in the muscle cells targeted by the larger 2°Mns (Wang and Brehm, 2017). The bases for these peripheral distinctions are explored in greater detail in the present study.

To address more directly the links between peripheral circuitry, Qc and behavior we have used direct imaging and *in vivo* paired recordings. This has led to identification of distinct calcium channel types mediating synaptic transmission in the two Mn types as well as a new organization involving sharing of synapses by both 1° and 2°Mn boutons. Moreover, the functional distinctions between 1° and 2°Mn boutons control two behaviors through shared synapses on the same muscle cell. This unexpected occupation of single synapses by functionally dichotomous boutons has not been reported in the nervous systems of any other vertebrate species.

## Results

### Innervation Pattern and Distribution of 1° and 2°Mn synapses

Individual fast muscle fibers receive innervation from a single 1°Mn and up to 3 additional 2°Mns (Westerfield et al., 1986). The distribution of synapses among fast muscle fibers is well documented for the 1°Mn types. The fast muscle is segregated into 4 quadrants within each segment, each bearing non-overlapping innervation by one of 4 different 1°Mns (Myers et al., 1986; Westerfield et al., 1986; Bello-Rojas et al., 2019). Our studies have focused on the CaP 1°Mn, which branches extensively to innervate all of the muscle cells in the ventral most quadrant (Fig. 1A). Labeling with either fluorescently conjugated alpha bungarotoxin (αBtx) or postsynaptic rapsyn, tagged with either GCaMP6f or GFP, reveals an en-passant distribution of postsynaptic acetylcholine receptor (AChR) clusters that number ~15 per muscle cell (Fig. 1A & B) (Wen et al., 2016; Brehm and Wen, 2019). Much evidence points to each cluster as representing an individual synapse formed with the 1°Mn. Morphologically, each of the ~15 receptor clusters co-localize with a fluorescence labeled 1°Mn terminal (Fig. 1B). Paired recordings and variance analysis of endplate currents (EPCs) also provided support for a one to one correspondence between 1°Mn terminals and receptor clusters (Wen et al., 2016). Direct test of the proposition that the 1°Mn innervates each individual receptor cluster formed on target muscle is now provided using an optical strategy (Fig. 1C & D). Transmitter release at individual synapses was monitored during CaP 1°Mn stimulation by the calcium indicator GCaMP6f, targeted to postsynaptic sites via fusion to rapsyn (Fig. 1B). To ensure that changes in fluorescence signal reflect calcium influx through the AChR during synaptic activity and not depolarization mediated release from intracellular stores, the measurements were made using the muscle dihydropyridine receptor mutant *relaxed* (Ono et al., 2001; Schredelseker et al., 2005). Owing to the high fractional calcium permeability of the AChR, our resolution was sufficient to detect single fusion events (Fig. 1C). Stimulation of the CaP 1°Mn at 0.2 Hz led to detectable GCaMP6f signals at every synapse on individual muscle cells (Fig. 1C & D). Through monitoring of the failures, this assay also allowed direct measurements of Pr at individual synapses (Fig. 1D & E). Consistent with the findings from paired recordings, Pr was near unity for each synapse (Wen et al., 2016). Collectively, these findings point to occupancy of nearly all synapses by the 1°Mn, raising the intriguing question as to the location of those synapses corresponding to the 2°Mns.

**Figure 1.**
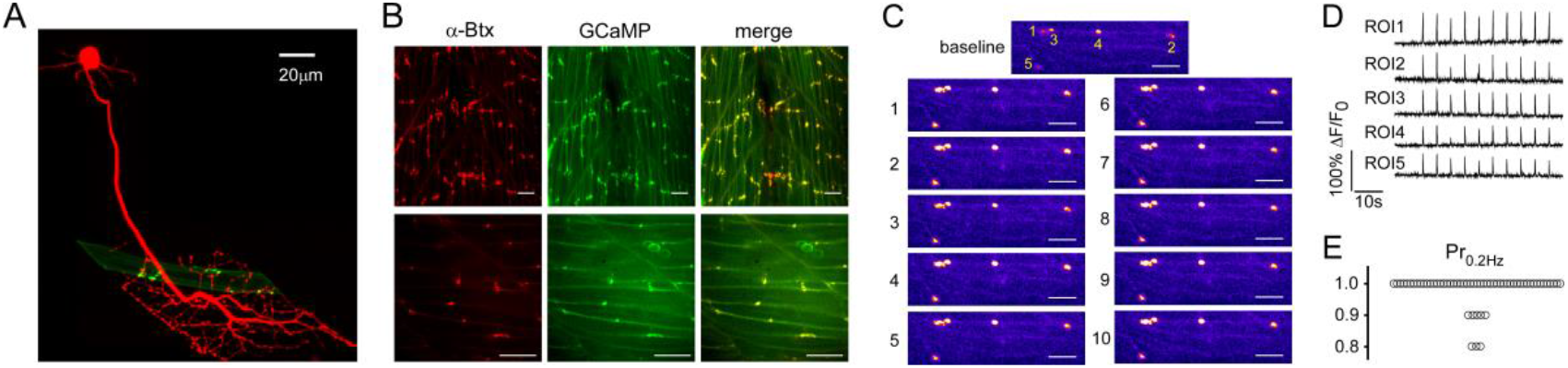
The CaP 1°Mn occupies nearly all of the synapses formed on individual muscle cells. (A) Maximal projection of confocal images of a CaP (red) and an example target muscle with its associated receptor clusters (green). The CaP was labeled with tdTomato expression driven by mnx1 promoter. Postsynaptic receptors were labeled with rapsyn-GFP driven by α-actin promoter that sparsely labeled muscle cells. (B) Confocal Z-stack (top panels) and single focal plane (bottom panels) images of ~2 segments showing co-localization of AChR labeling with a-Btx (red) and rapsyn-GCaMP6f basal fluorescence (green). Scale bars 20 μm. (C) Example fluorescence responses for 5 synapses in response to 0.2 Hz stimulation of the CaP (10 consecutive responses shown). In this focal plane, ROIs 1-4 are located in the same muscle cell, while ROI 5 is on a different muscle. (D) Individual GCaMP6f responses for the 5 ROIs in response to 0.2 Hz stimulation of the CaP. (E) Overall distribution of Pr at 0.2 Hz measured for 59 ROIs from 16 fish.

We sparsely labeled 2°Mns with transient expression of fluorescent proteins to facilitate reconstruction of branching patterns for individual 2°Mns. Consistent with a recent publication we identified a broad range of branching patterns, from simple to highly complex, among the over 40 2°Mns (Bello-Rojas et al., 2019)(Fig. 2A). Both the extent of branching and synapse numbers was directly related to soma size (Fig. 2A). These findings indicate that larger 2°Mns innervate a larger number of muscle cells, accounting in part for the size dependence of power production.

**Figure 2.**
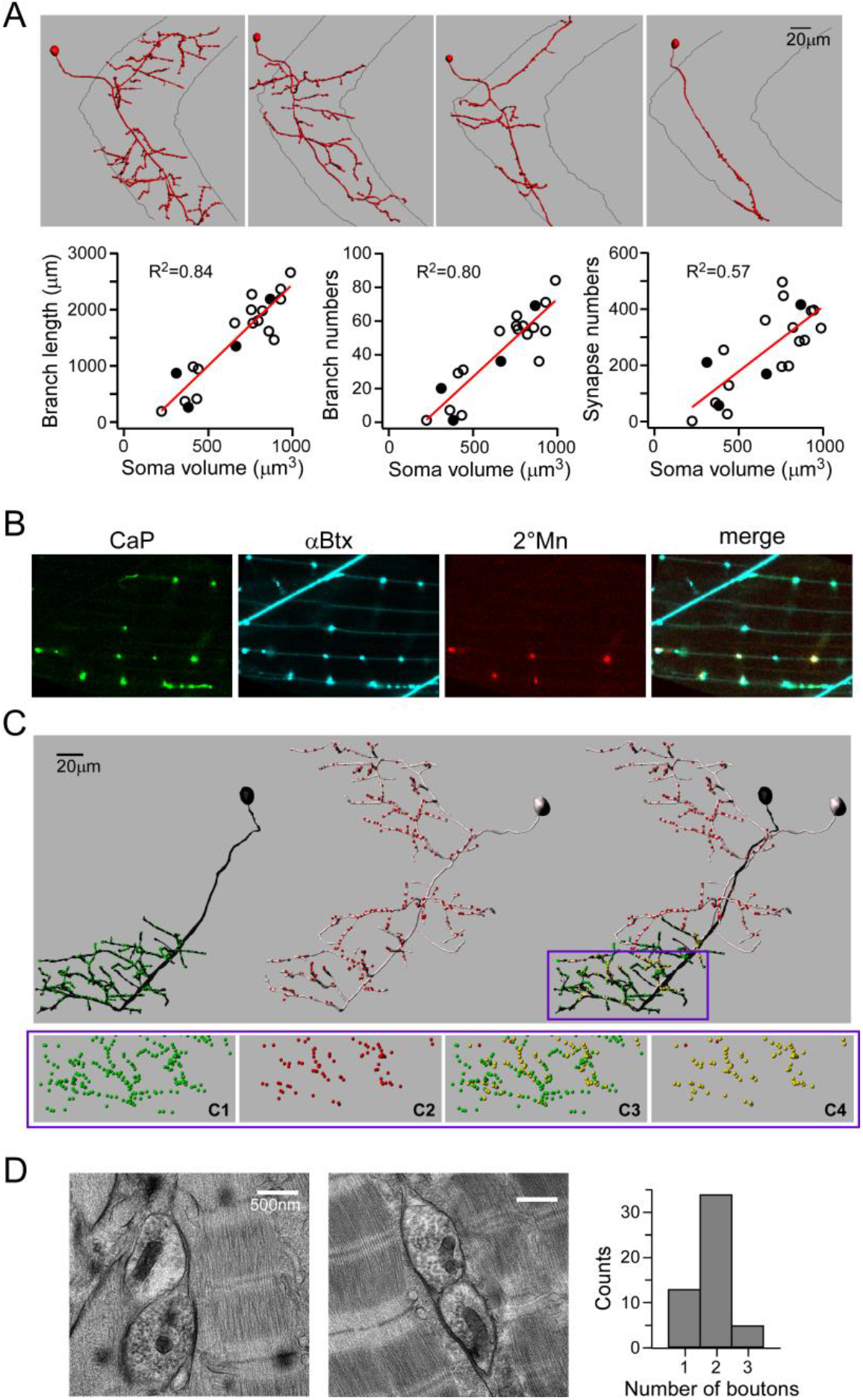
1° and 2°Mns share synapses. (A) Imaris reconstructions of four different 2°Mns exemplifying the range of branching patterns (top panels). Relationships between soma volume versus axonal branch length, branch number and αBtx puncta number are shown (bottom panels). Each symbol represents a reconstruction of an individual 2 Mn with filled symbols corresponding to the 4 examples shown. The relationship shows a linear fit with R^2^ values indicated. (B) Confocal images showing co-localization of fluorescently labeled CaP terminals (green), postsynaptic AChR (cyan), 2°Mn terminals (red) and merge. (C) Imaris reconstruction of a 1°Mn (black) and a 2°Mn (gray). Individual synapses, based on aBtx label, are color-coded in green (for 1°Mn), red (for 2°Mn) and yellow (scored as shared). Only the bottom half of the ventral muscle field is considered for quantitation as CaP is the sole 1°Mn innervating that area. An expanded view of the boxed region below shows the distribution of synapses that are associated with CaP (C1), 2°Mn (C2), scored as shared among all synapses (C3) or in relation to the 2° Mn synapses (C4). In this example 98% of the labeled 2°Mn synapses in the overlapping region were shared with the CaP. (D) Example electron micrographs showing two boutons sharing a synapse. Histogram of the 52 images from 6 fish is shown (right).

To examine the location of 2°Mn synapses relative to those of the 1°Mn, sparse labeling of 2°Mns was performed in a transgenic line expressing EGFP specifically in CaP 1°Mn. Single 2°Mns that branch into ventral target territory of the CaP 1°Mn were imaged along with the CaP arbor and aBtx labeled AChR clusters (Fig. 2B & C). Coincidence labeling of AChR clusters with the terminals of both the CaP 1°Mn and 2°Mn pointed to frequent sharing of synapses between the two Mn types (Fig. 2B). To quantitate the extent of sharing 3D reconstruction was performed for the synapses in the CaP target territory for both Mn types (Fig. 2C). As expected, over 99% of all synapses in this region were associated with the CaP (Fig. 2C1; 99.4 +/− 0.5%, n=13). In addition, approximately 98% of the receptor clusters associated with the labeled 2°Mn (Fig. 2C2) co-localized with those of the CaP (Fig. 2C3 & 2C4; 97.9 +/− 1.6%, n=13). Because this experiment labeled only one of the 2°Mns that extended branches into the CaP territory, those CaP synapses that were scored as not shared (Fig. 2C3) are either shared with a different unlabeled 2°Mn and/or occupied solely by the CaP.

This method did not provide information as to how many 2°Mns share individual synapses with a 1°Mn. For this purpose we turned to conventional electron microscopy to count the bouton number at individual synapses. To satisfy the criteria for a synapse we required that there be evidence of electron dense material between muscle and nerve as well as accumulation of synaptic vesicles (Fig. 2D). The overall distribution determined for 6 fish ranged from 1 to 3 boutons (n= 52 synapses) (Fig. 2D). Assuming that in all cases one of the boutons corresponds to the 1°Mn, we interpret this data as representing between 0 and 2 boutons of 2°Mns at each synapse, with a major mode of 1. It is likely that these counts represent an underestimate due to missed Mn boutons out of registry during sectioning.

### Functional distinctions between 1° and 2°Mn mediated neuromuscular transmission

Paired recordings between target muscle and either the CaP 1°Mn or a 2°Mn revealed Mn specific differences in neuromuscular transmission (Fig. 3). As shown by use of optical recordings (Fig. 1) and published paired recordings, the CaP 1°Mn has near unity Pr at low stimulus frequency. As such it represents a fast depressing synapse when the stimulus frequency is increased (Wen et al., 2016). It also has a Qc of ~15 that matches with estimated number of functional release sites and the number of postsynaptic receptor clusters (Fig. 3A) (Wen et al., 2016). By contrast, the 2°Mn has smaller and highly variable Qc (Fig. 3A & C) (Wang and Brehm, 2017). For quantitative comparison of synaptic strength to the CaP 1°Mn, we chose a specific subgroup of 2°Mns that extends branches into both dorsal and ventral muscle segments with an input resistance in the range of 250-500 MΩ (one example shown in first panel of Fig. 2A)(Bello-Rojas et al., 2019). EPC amplitudes for the 2°Mn showed large fluctuations along with frequent failures (34 +/− 27% failure rate, n=21; Fig. 3B). These fluctuations in amplitude translate directly to differences in Qc based on similar mean unitary (mEPC) amplitudes for both Mn types (Fig. 3C). The mEPC amplitude was determined for each Mn using isolated unitary events that are not synchronous with action potential firing, taking advantage of the fact that high frequency stimulation led to an elevated occurrence of asynchronous mEPCs (Wen et al., 2010; Wen et al., 2016). The mean amplitudes corresponded to 945 +/− 216 pA (n=20) for the 1°Mn and 926 +/− 155 pA (n=14) for 2°Mn. Based on the direct method (mean EPC amplitude/mean mEPC amplitude) the Qc averaged 1.2 +/− 1.0 (n=21) for the 2°Mn, and 12+/− 3.2 (n=26) for the CaP 1°Mn. The difference in synaptic strength between Mn types reflects the much lower Pr in 2°Mns compared to 1°Mn synapses. The Pr in 2°Mn can be further facilitated during repetitive stimulation, resulting in reduced occurrence of failure and increased EPC amplitudes (average fold of increase 2.1 +/− 0.6, n=12; Fig. 3D).

**Figure 3.**
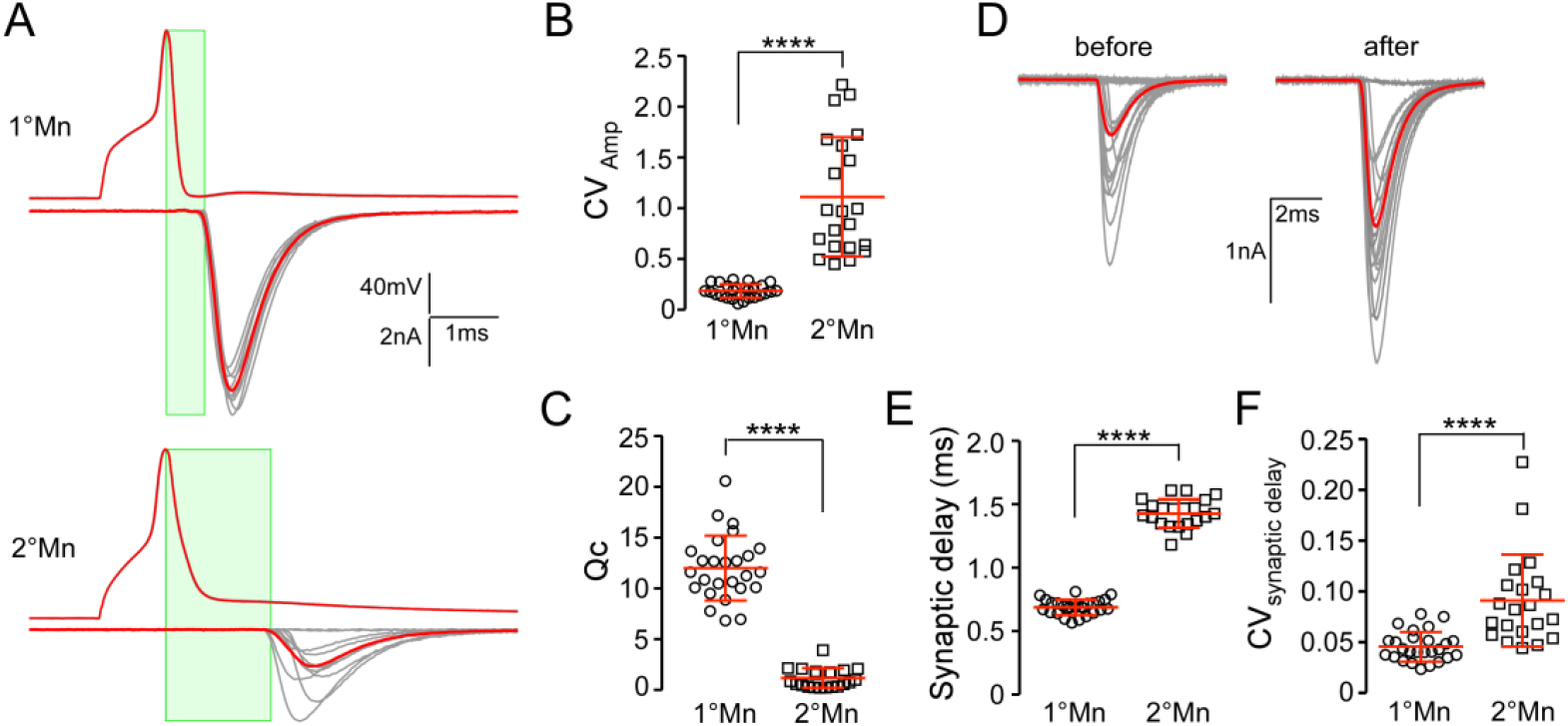
Functional distinctions between 1° and 2°Mn synapses. A. Example traces of action potentials and EPCs (average in red) for CaP 1°Mn and 2°Mns in response to 10 stimuli at 1 Hz. Time window for synaptic delay measurement is shown (in green). B. Scatter plot comparing coefficient of variation (CV; mean +/− S.D. in red) for EPC amplitudes between the 1° and 2°Mn. CV is calculated for 20 responses acquired at 1 Hz for each cell. C. Scatter plot comparing quantal content (Qc; mean +/− S.D. in red) between 1° and 2°Mns. D. A sample 2°Mn paired recording showing 20 EPCs at 1 Hz stimulation before and after facilitation (average in red). The facilitation was induced with a 100 Hz, 2 s stimulus train in the 2°Mn. E. Scatter plot comparing the synaptic delay (mean +/− S.D. in red) between 1° and 2°Mn. F. Scatter plot comparing CV (mean +/− S.D. in red) for the synaptic delay between 1° and 2°Mn. **** indicates P<0.0001 in panel B, C, E and F.

The EPCs for the two Mn types have similar rise and decay times but the onset of the EPC is delayed in 2°Mn when compared to the 1°Mn. Measured as the interval between the peak of the Mn action potential and 10% rise of EPC, the synaptic delay averages 1.43 +/− 0.11 (n=21) for the 2°Mn compared to 0.69 +/− 0.06 (n=26) for the 1°Mn, corresponding to a 2.1-fold difference (Fig. 3E). Additionally, there are greater variations in the synaptic delay for the 2°Mn EPCs (Fig. 3F). Overall, 2°Mn synapses are weaker in strength, less reliable and lack the temporal precision of 1°Mn synapses.

### Synaptic transmission and behavior in a P/Q calcium channel null mutant

Assignment of P/Q calcium channels as mediators of 1°Mn neuromuscular transmission were based on a mutant line harboring a missense mutation in the P/Q channel (Wen et al., 2013). This mutant showed greatly reduced levels of neuromuscular transmission *in vivo* and compromised swimming, both of which we attributed to the residual levels of P/Q expression (Wen et al., 2013). We have subsequently obtained a second P/Q mutant line that represents a complete null due to premature truncation of the channel and this has provided insights into functional distinctions among Mn types. The first hints came from monitoring of electrical stimulation induced escape and swimming using high speed imaging (Fig. 4). Wild type fish responded to a brief shock with an initial powerful C-bend characterized by a high degree of curvature corresponding to 131 +/− 22 degrees (Fig. 4A & B; n=12 fish). This bend was followed by a bout of rhythmic swimming with alternating tail-beats that decreased in strength and duration. In the P/Q null fish the initial bend reached only ~1/3 of the curvature measured for C-bend in wild type fish (49 +/− 15 degree, n=14; Fig. 4A & B). Additionally the onset of the first body bend was delayed compared to wild type fish (Fig. 4A & C; 21.0 +/− 4.5 ms, n=12 for wt; 39.4 +/− 5.4 ms, n=14 for P/Q null). This delay of 18.4 ms compared favorably to the average duration of the C bend for wild type fish (19.1+/− 6.4 ms, n=12). These measurements demonstrated that P/Q null mutants lack C-bend while maintaining the ability to mount rhythmic swimming.

**Figure 4.**
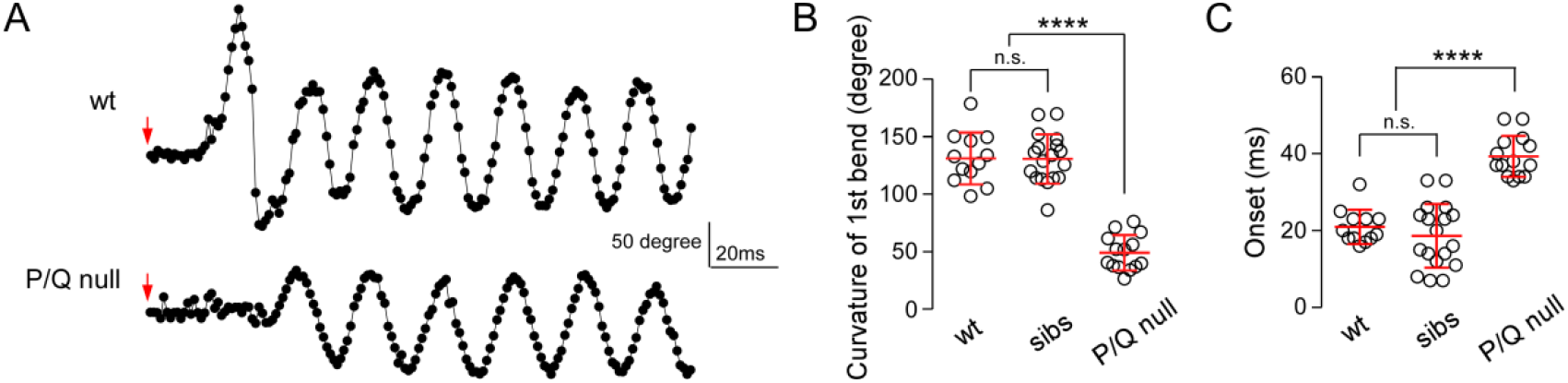
P/Q-type mediated transmission specifically mediates the escape response. A. Example time series showing changes in body curvature during responses triggered by a 20 ms electrical field stimulation (arrows are the start marks) of a wild type and P/Q null fish. Movies were acquired at 1000 frame/s and analyzed using Flote motion analysis software. B. Scatter plot comparing curvature of the first bend for wild type, siblings and P/Q null mutants. C. Scatter plot comparing the onset of movement for wild type, siblings and P/Q null mutants. Heterozygous siblings were indistinguishable from wild type. n.s., not significant with p values greater than 0.05. Mean+/− S.D. is indicated in red. ****p<0.0001.

Using paired recordings, we revisited previously published evidence pointing to reliance on P/Q type calcium channels in neuromuscular transmission between the 1°Mn and target muscle. The total charge transfer for 20 responses at 1 Hz corresponded to 0.55 +/− 0.59 pC (n=17) in the P/Q null mutants, compared to 204 +/− 71 pC (n=29) for wild type fish, reflecting a >99% reduction of synchronous release in the mutant (Fig. 5A & B).

**Figure 5.**
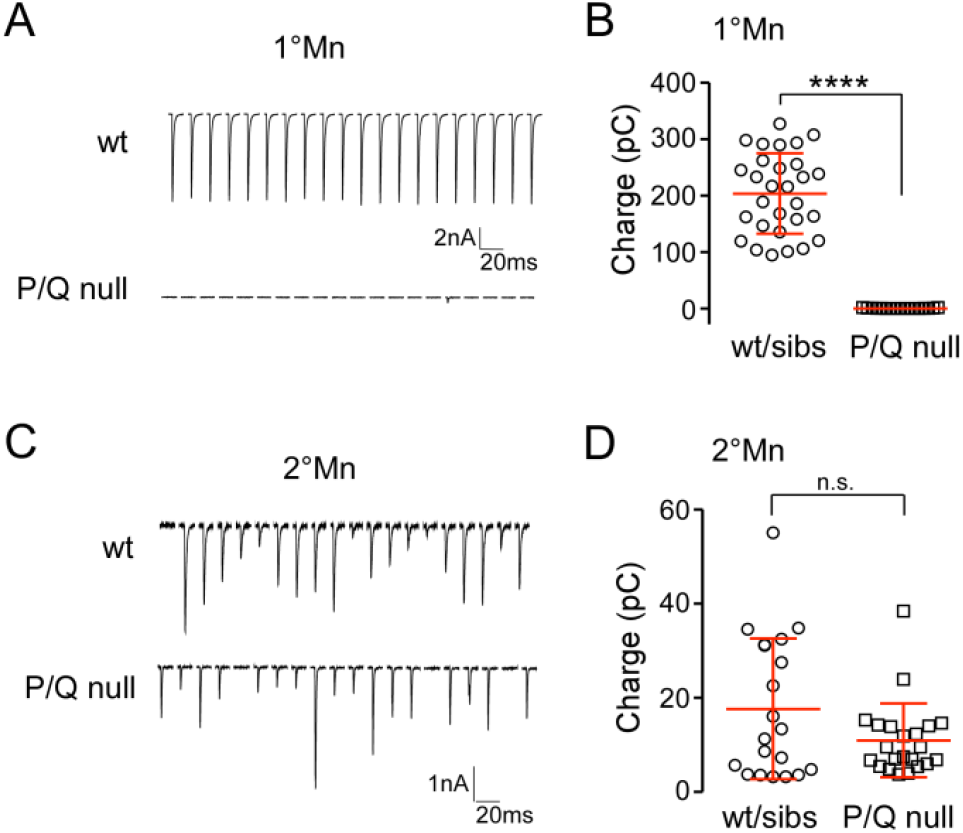
The P/Q channel null mutant specifically abolished synaptic transmission in 1°Mns. A. Example traces of 20 consecutive EPC responses at 1 Hz in CaP 1°Mn from a wild type (top) and a P/Q null (bottom). B. Scatter plot comparing the total charge transfer for 20 responses at 1 Hz in CaP 1°Mn between wild type and P/Q null mutants. Mean+/− S.D. is indicated in red. ****p<0.0001. C. Example traces of 20 consecutive EPC responses at 1 Hz in a 2°Mn from a wild type (top) and a P/Q null (bottom). D. Scatter plot comparing the total charge transfer for 20 responses at 1 Hz in wild type and P/Q null mutants. Mean+/− S.D. indicated in red. n.s. corresponds to p=0.07. Heterozygous siblings were indistinguishable from wild type fish, and data were pooled in the summary plots.

As 2°Mns have been shown to be central to rhythmic swimming we performed paired recordings between 2°Mns and target muscle in the P/Q null mutants. In contrast to the absence of release in the 1°Mns, evoked synaptic transmission was intact between 2°Mn and target muscle (Fig. 5C & D). Comparison of total release showed no statistically significant difference between wild type and P/Q null fish (Fig. 5D; 17.7 +/15 pC, n=20 for wt and sibling; 11.0 +/− 7.8, n= 22 for mutants. P=0.07). Overall, our electrophysiology and behavior measurements indicate that a calcium channel other than the P/Q type mediates neuromuscular transmission in the 2°Mns.

### Identifying the calcium channel type used for synaptic transmission in 2°Mns

The zebrafish P/Q channel type has a pharmacological profile that sets it apart from its mammalian counterpart (Wen et al., 2013). Specifically, zebrafish P/Q channels are sensitive to ω-conotoxin GVIA, a toxin widely considered to be N-type specific. ω-conotoxin GVIA blocked both zebrafish P/Q type and N type currents in heterologous expression system as well as abolishing CaP 1°Mn synaptic transmission *in vivo* (Wen et al., 2013). To test the effect of the toxin on 2°Mn mediated synaptic transmission we used a line of fish expressing channelrhodopsin (ChR-YFP) principally in the 2°Mns (Wyart et al., 2009). Recordings revealed that application of 1 μM ω-conotoxin GVIA blocked all the light-evoked responses (n=4). Additionally, injection of ω-conotoxin GVIA into fish leads to total paralysis of the animal (n=15). On the basis of sensitivity to ω-conotoxin GVIA and having ruled out the P/Q type, we tested for N type calcium channel involvement in 2°Mn mediated neuromuscular transmission.

To date, efforts to identify or generate zebrafish N and P/Q subtype specific antibodies have failed, due largely to the high sequence similarity between the channel types. As an alternative means we turned to a panel of 9 ω-conotoxins that have been tested on mammals to be N-type specific (Alomone labs) in search for a toxin with differential blocking effect on zebrafish N and P/Q channels. The toxins were first tested for their effectiveness on blocking 2°Mn mediated synaptic transmission. For this purpose high concentrations (1-5 μM) of each toxin were applied to fish expressing ChR-YFP in 2°Mns, and light evoked responses were recorded in the muscle. Four toxins showed no evidence of block and were not subject to further testing (ω-conotoxin CVIA, SVIB, SO3, and FVIA; Table 1). The remaining 5 toxins strongly blocked of 2°Mn transmission and were then tested for specificity in paired recordings between the CaP 1°Mn and target muscle. Application of ω–conotoxins RVIA and MVIIA also blocked CaP mediated transmission indicating that they act on both the zebrafish P/Q and N type channels and were not further tested. By contrast, ω–conotoxins CVIE, CVIF and CnVIIA showed greatly reduced sensitivity on the CaP 1°Mn release compared to their effects on 2°Mn mediated transmission. Consequently, they served as candidates for specific block of the putative N type channel responsible for 2°Mn transmission (Table 1). Two of these were used for further studies.

**Table 1.** Exploratory screen for toxins that distinguish between zebrafish N-type and P/Q type calcium channels

| Toxins | ChR-YFP response | CaP release | likely outcome |
| --- | --- | --- | --- |
| ω-conotoxin CVIA | not blocking at 1 μM | N.A. | neither N nor P/Q blocking |
| ω-conotoxin SVIB* | not blocking at 1 μM | not blocking at 1 μM | neither N nor P/Q blocking |
| ω-conotoxin SO3 | not blocking at 1 μM | N.A. | neither N nor P/Q blocking |
| ω-conotoxin FVIA | not blocking at 1 μM | N.A. | neither N nor P/Q blocking |
| ω-conotoxin RVIA | strong block at 2 μM | strong block at 2.6 μM | both N- and P/Q blocking |
| ω-conotoxin MVIIA | strong block at 1 μM | nearly complete block at 1 μM | both N- and P/Q blocking |
| ω-conotoxin CVIE | strong block at 1 μM | slight block at 330nM | N-specific |
| ω-conotoxin CVIF | strong block at 1 μM | residual response at 1 μM, 500nM, 330nM | N-specific |
| ω-conotoxin CnVIIA | strong block at 5 μM | slight block at 2.5 μM | N-specific |

\*When tested subsequently on the basis of behavior $\omega$ -Conotoxin SVIB caused paralysis indicating it has actions beyond calcium channels, likely directly on muscle.

To confirm a preferential block of the N type, we quantitated the potency of the toxins on heterologously expressed zebrafish N and P/Q type channels (Fig. 6). Both calcium channel types were expressed in HEK293T cells as previously published (Wen et al., 2013). For each toxin at least 4 concentrations were tested for block of calcium current. The half maximal inhibitory concentration, IC_50_, were compared between the two channel types (Fig. 6A). The higher affinity CVIF showed a 6.8 fold preference for block of N type with an IC_50_ corresponding to 44 nM for N type and 301 nM for P/Q type (Fig. 6B). The lower affinity blocker CnVIIA showed a 2.7 fold difference with the IC_50_ corresponding to 904 nM for N type and 2.4 μM for P/Q type (Fig. 6C).

**Figure 6.**
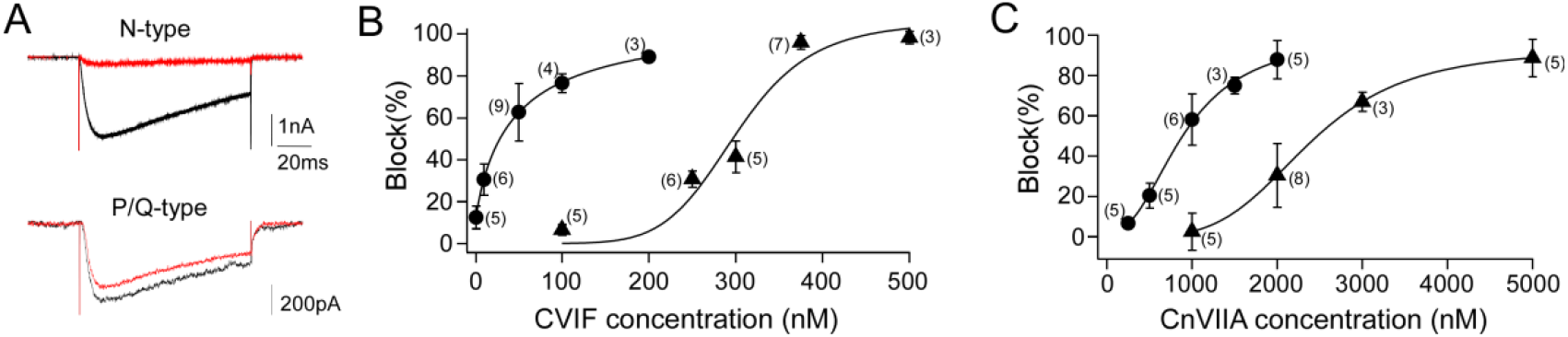
Two ω-conotoxins that distinguish zebrafish N and P/Q type calcium channels. A. Sample recordings of calcium currents from HEK293T cells expressing zebrafish P/Q or N type channels, before (black) and after (red) application of ω-conotoxin CVIF. 200nM and 250nM of toxin were applied for N and P/Q, respectively. B. Dose response curve of ω-conotoxin CVIF block on N (circles) and P/Q (triangles) currents. C. Dose response curve of ω-conotoxin CnVIIA block on N (circles) and P/Q (triangles) currents. For the dose responsive curve, each symbol represents the average of multiple applications for the indicated number of different cells tested. The average values for all cells tested at each concentration are shown along with S.D. Each relationship was fit by a Hill equation.

With the concentration dependencies for N versus P/Q channels in hand we turned to *in vivo* paired recordings to test for selective actions on 2° and 1°Mns mediated synaptic transmission (Fig. 7). For this purpose ω–conotoxin CVIF was selected because it provided the greatest distinction between the channel types heterologously expressed in HEK cells. At 200 nM CVIF blocked ~90% of N type and less than 20% P/Q type calcium current in HEK cells (Fig. 6A & B). EPCs were recorded for each Mn type before and after a solution change to 200 nM CVIF. The blocking actions mirrored those obtained from expressed channels with an average block of 21.7+/− 7.6% (n=5) for the CaP 1°Mn and 73.5+/− 8.5%(n=5) for the 2°Mn (Fig. 7A & B). Exacting correspondences between the measured channel block in HEK293 cells and effects on release of transmitter *in vivo* would not be expected in light of the non-linear relationship between calcium entry and transmitter release as well as other uncontrolled factors such as the calcium channel subunit composition. Nevertheless, the clear differential action of the toxin on the two Mn types provided strong support for the key role of N-type channel in 2°Mn mediated neuromuscular transmission.

**Figure 7.**
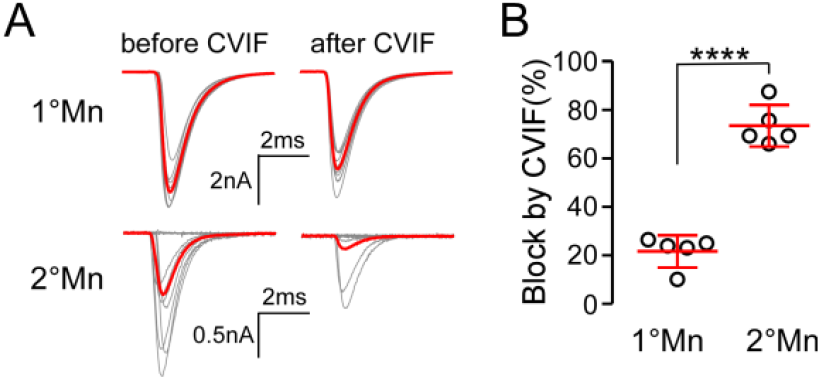
N-type specific ω-conotoxin CVIF preferentially blocks transmitter release in 2°Mn. A. Example paired recordings showing block of EPCs by CVIF in 1° and 2°Mns. 10 EPCs along with the average (in red) at 1 Hz stimulation are shown before and after application of 200 nM toxin. Decreased responses in 2°Mn was accompanied by an increased failure rate, with the smallest responses representing miniature EPCs. B. Scatter plot comparing CVIF block in 1° and 2°Mn EPC recordings. Synaptic responses were quantified as the summed charge transfer of 20 EPCs. ****p<0.0001.

## Discussion

When first published, our identification of a zebrafish P/Q calcium channel as the mediator of neuromuscular transmission came as a surprise (Wen et al., 2013). Frog NMJ had long been associated with reliance on an N type calcium channel based on sensitivity to ω-conotoxin GVIA and MVIIC, setting it apart from mammalian NMJs that used a P/Q type (Sano et al., 1987; Uchitel et al., 1992; Katz et al., 1995; Thaler et al., 2001). Thus, it was assumed that higher and lower vertebrates differed along this basic standpoint. However, this distinction has always been difficult to reconcile in light of such an otherwise highly conserved synapse among vertebrates. Pharmacological testing of ω-conotoxin GVIA, presumed to be N type specific, on heterologously expressed zebrafish P/Q and N type calcium channels provided a potential resolution (Wen et al., 2013). Both channel types showed similar blocking profile, pointing to a potential mis-identification in frog and to common usage of a P/Q type among all vertebrates.

Subsequent paired recordings from CaP 1°Mn and target muscle from a P/Q channel mutant lend strong support for the reliance of zebrafish neuromuscular transmission on this channel type (Wen et al., 2013). One caveat, however, is reflected in the ability of the P/Q mutant line to retain limited motility. At the time it was reasoned that the mutant line was not a complete null. The newly obtained P/Q mutant line in the present study was able to mount rhythmic swimming, despite the prediction that it represents a functional null. This finding, together with the observation that all swimming movement was completely blocked by ω-conotoxin GVIA, raised the likelihood of an N type calcium channel as mediator of release in 2°Mn, the Mn type though to support rhythmic swimming. As a test of this proposition we identified two conotoxins that preferentially blocked zebrafish N type channels over P/Q types expressed in HEK cells. *In vivo* testing using paired recordings then confirmed a preferential block of 2°Mn over 1°Mn synaptic transmission. On these bases we conclude that 1°Mns use a P/Q type channel and the 2°Mns use an N type channel for neuromuscular transmission in zebrafish. These toxins may now provide a means for further testing roles for P/Q versus N type calcium channels at the frog NMJ.

Our finding that the P/Q null mutants lacked a C-bend is consistent with published studies pointing to specific dependence on the 1°Mns for this behavior (Liu and Westerfield, 1988; Fetcho and O’Malley, 1995). The rhythmic swimming still occurred in the mutant, indicating that the 2°Mns do not participate in the C-bend. The behavioral responses associated with 1° and 2°Mn types also match well with differences in Qc between Mn types and associated differences in contraction strength. This is due in part to the peripheral circuitry where small sized 2°Mns contact fewer muscle cells thereby generating less power and slower swimming (Fig. 2A)(Bello-Rojas et al., 2019). In addition, the smaller Mns also have smaller Qc, leading to the generation of weaker contractions at the level of individual muscle cells (Wang and Brehm, 2017). The small zebrafish muscle cells are nearly isopotential due to a high input resistance, thereby distributing the synaptic depolarization throughout the cell. The 1°Mn produce the largest depolarization due to the large Qc which helps insure that the C-bend generates the most power and the fast depression insures that only a single such response is generated. By contrast the 2°Mns vary greatly in their Qc, accounting for a graded depolarization and release of intracellular calcium in muscle (Wang and Brehm, 2017). This enables graded power development for rhythmic swimming that span a range of speeds.

What role might the different calcium channel types play in conferring Mn specific function? The functional distinctions center principally on the strong transmission with a high Pr in P/Q type mediated 1°Mn versus weak transmission with a low Pr in N type mediated 2°Mn. These distinctions could arise from difference in the intrinsic biophysical properties of the channels. At zebrafish NMJ and mouse mossy fiber synapses, the P/Q type has been shown to have a higher open probability than N type, which would be expected to result in greater calcium influx upon action potential (Li et al., 2007; Naranjo et al., 2015). In addition, calcium accumulation also depends on both the density and location of calcium channels (Dittrich et al., 2018; Rebola et al., 2019). Should the P/Q channels occur at higher density the domains formed by calcium influx would experience greater overlap, thereby increasing local calcium concentration (Stanley, 2016). Furthermore, there might be channel type specific differences in coupling between presynaptic calcium influx and the calcium sensor for release, a critical determinant for synaptic efficacy (Eggermann et al., 2011; Rebola et al., 2019). Tighter coupling has been shown for the several central synapses that rely on P/Q type, while looser coupling is often associated with weaker synapses that use the N-type for release (Iwasaki and Takahashi, 1998; Wu et al., 1999; Forti et al., 2000; Stephens et al., 2001; Hefft and Jonas, 2005; Bucurenciu et al., 2010). Consistent with this idea, we observed a longer and more variable synaptic delay for 2°Mn synapses. The P/Q and N-type channels have been shown to differentially interact with the multiple proteins in the exocytotic machinery and other active zone components (Mochida et al., 2003; Cao and Tsien, 2010; Davydova et al., 2014), providing a potential molecular mechanism for the difference in coupling.

Innervation of individual muscle cells by functionally dichotomous Mns, as reported here, has also been identified at the larval wall muscles in Drosophila (Newman et al., 2017). There, the 1s and 1b Mns innervate the same muscle cells but differ in terms of reliability and fatigue. The 1s synapses are reliable and depress compared to 1b synapses that are often largely silent and ‘wake up’ in response to high frequency stimulation (Newman et al., 2017). Thus, the 1s is similar to the 1°Mn type and the 1b to 2°Mns that show frequent failure. The functional distinctions at the fly NMJ have been shown to involve postsynaptic glutamate receptor composition, and presynaptic differences in calcium channel density as well as the levels of the active zone organizer *bruchpilot* (Akbergenova et al., 2018). At zebrafish NMJ we can rule out involvement of postsynaptic receptors in the distinction because of synapse sharing. Zebrafish NMJ offers some significant advantages over flies in the hunt for mechanisms regulating Pr. Simplicity is provided by the low number of synapses per muscle cell (~15 vs hundreds in fly), the large size of individual synapses which precludes need for super resolution methodology, and the amenability to paired patch clamp Mn-muscle recording (Brehm and Wen, 2019). Additionally, sharing of 1° and 2° Mn boutons at the same synapse offers the opportunity for side by side interrogation of high and low Pr synapses for differences at the molecular and ultrastructural level. High pressure freeze electron microscope tomography of zebrafish NMJs for example, will offer the opportunity to test for differences in presynaptic organization between the two Mn types (Helmprobst et al., 2015).

The sharing of individual neuromuscular synapses by multiple Mns is not without precedent among the vertebrates. During early development and re-innervation following injury, multiple different Mns share the same synaptic gutter and receptors (Brown et al., 1976). Through a process of competition, Mns sequentially withdraw over the course of development until only one Mn terminal remains at every synapse (Grinnell, 1995). The mechanisms causal to the competition remain uncertain but evidence points to differences in synaptic strength (Jordan et al., 1992; Barber and Lichtman, 1999; Turney and Lichtman, 2012). Our data from zebrafish NMJs showing a preponderance of a single 2°Mn co-occupying each synapse along with the 1°Mn now points to a competition among 2°Mns. This would indicate that whatever mechanisms are responsible for competition within a class of Mn, does not apply to 1°Mns versus 2°Mns. This could potentially play a central role in establishment of stereotypic peripheral circuitry through timed outgrowth and competition. According to this model the axons of the 1°Mns, which are the first to emerge from the spinal cord, are followed by timed outgrowth of the numerous 2°Mns. We propose that the first 2°Mns to emerge would be the most branched and co-occupy the greatest number of synapses due to availability. 2°Mns emerging later would co-occupy fewer synapses due to prior occupancy by another 2°Mn. In this manner a stereotypic innervation pattern could potentially be formed through precisely timed outgrowth of 2°Mns, with those bearing the fewest branches and synapses reflecting the last to grow out.

Collectively, this study points to distinct roles played by P/Q versus N type calcium channel in mediation of two specific behaviors. It does so through use of a newly described peripheral circuitry involving synapse sharing by functionally distinct boutons, one low Pr and the other high Pr. This sharing endows each postsynaptic muscle the ability to carry out the specialized behavior, a process generally associated with activation of different muscle cells in most vertebrate classes. Prior to this study the sharing of individual excitatory synapses as a process of circuitry organization has not been reported, calling for further investigation at CNS synapses.

## Material and Methods

### Fish lines

All zebrafish (*Danio rerio*) were maintained in the in-house facility. The P/Q null mutant fish line (cacna1ab~sa35120) was obtained from Sanger Institute’s Zebrafish Mutation Project (Kettleborough et al., 2013)(Zebrafish International Resource Center, Eugene, Oregon). It harbors a nonsense mutation that truncates the protein at amino acid 372 (out of a full length of 2280). Zebrafish husbandry and procedures were carried out according the standards approved by Institutional Animal Care and Use Committee at Oregon Health & Science University. All experiments were performed using larva at 72-120 hours post fertilization (hpf). The sex of the larva was unknown at this age.

### Motoneuron labeling and reconstruction

A double transgenic line SAIGFF213A; UAS:GFP provided the labeling of CaP 1°Mn. It expresses EGFP in a subset of spinal neurons, exclusively in CaP in some segments (Muto et al., 2011). For sparse labeling of 2°Mn, we transiently expressed mCherry-tagged synaptotagmin 2 driven by the neuronal specific HuC promoter by injecting the plasmid into single cell embryos. In some experiments Gal-UAS system was used to achieve mosaic expression by co-injection of two DNA plasmids. In those cases mnx1 drove the expression of Gal4 in motoneurons (mnx1:Gal4 plasmid, kindly provided by Dr. David Mclean, Northwestern University) and a red fluorescent protein containing UAS construct served as the reporter (UAS:tdTomato; or UAS:Cytobow plasmid, kindly provided by Dr. Tamily Weissman-Unni, Lewis & Clark College). Injected wild type fish were screened at 72-96 hpf for single fluorescent 2°Mns that could provide detailed morphology. For experiments investigating synapse sharing between Mn types, injected SAIGFF213A; UAS: GFP larva were screened for candidates containing segments where a single 2°Mn shared an overlapping target field with the CaP 1°Mn. Candidate fish were subsequently injected in the aorta with 0.5 nL of 1mg/ml CF405s conjugated a-Btx (Biotium, Fremont, California) and allowed 15-30 minutes to label AChR clusters before mounted for imaging.

Z stack images of labeled Mns and a-Btx labeled AChR clusters were acquired using a laser scanning confocal microscope (Zeiss LSM 710) with a C-Apochromat 40x/ 1.2W objective. The z stack images of soma and axon branches were reconstructed using Imaris 9.0 filament software (Bitplane, Zurich, Switzerland). a-Btx punta were detected using the Spots function and annotated as spheres of 2 μm radius. To associate a postsynaptic receptor cluster with either Mn type, a maximum of 1μm distance between a neuronal branch and individual a-Btx puncta was used as the criteria.

### Optical monitoring of neuromuscular transmission at single synapses with GCaMP6f

Our transgenic line targeting the genetically encoded calcium sensor, GCaMP6f (Chen et al., 2013), to postsynaptic receptor clusters was used to monitor neurotransmitter release at individual single synapses. GCaMP6f was fused to the C-terminal of AChR clustering protein rapsyn, and fusion protein expression driven with muscle specific α-actin promoter. To ensure the fluorescence change is not complicated by calcium release from intracellular store, the rapsyn-GCaMP6f fish was crossed with the *relaxed* mutant that lacks functional dihydropyridine receptors (Ono et al., 2001; Schredelseker et al., 2005). Paralytic *relaxed* homozygous fish were used for GCaMP6f monitoring.

Fluorescence response during 0.2 Hz stimulation of the CaP was monitored using CSU10 spinning confocal microscope (Yokogawa Electric, Japan) configured with ImagEM X2 EM-CCD camera (Hamamatsu Photonics, Japan). Images were acquired at a rate of 30 frames per second using Micro-Manager Software (UCSF, San Francisco, CA). Frame by frame analysis of fluorescence responses was performed in ImageJ (NIH, Bethesda, MD). Individual areas of interest (ROIs) were defined based on the basal fluorescence of GCaMP6f that provided the labeling of synapses. ΔF/F_0_ was determined for each ROI, where F0 represents the pre-stimulus baseline obtained by averaging 1 s of pre-stimulus images, and ΔF the difference between fluorescence signal and the baseline (F-F0). Positive responses were identified as peaks greater than 10 times of the standard deviation of the baseline signal. Pr was determined as the ratio of the number of positive responses to the total number of stimuli.

### Electrophysiology of neuromuscular transmission

Paired recordings between Mns and target fast skeletal muscle were performed as previously described for CaP and its muscle cell target (Wen and Brehm, 2010). CaPs were identified on the basis of stereotypical morphology and biophysical properties. In light of the large diversity among 2° Mns, we focused on the subset extending branches to both ventral and dorsal muscle targets) for quantitation. To identify 2° Mns for recording, we relied on sparse labeling with mnx1:Gal4 and UAS:tdTomato plasmid injection into single cell embryos or dye filling with 40 μM Alexa Fluor 568 hydrazide in the neuronal recording pipette. The recording bath solution contained (in mM): 134 NaCl, 2.9 KCl, 1.2 MgCl2, 2.1 CaCl2 and 10 Na-HEPES, pH 7.2, ~290 mOsm. Both neuron and muscle recording pipettes contained (in mM: 115 K gluconate, 15 KCl, 2 MgCl2, 5 K-EGTA, 10 K-HEPES, 4 Mg-ATP, pH 7.2, ~ 290 mOsm). Action potentials in Mns were elicited by 1 ms current injection, while EPCs were recorded from target muscles voltage clamped at −50 mV. Muscle recording pipettes had a resistance of 2-5 MOhm and series resistance was kept under 10 MΩ. All recordings were performed with 60% online series resistance compensation. Synaptic currents were sampled at 100 kHz and filtered at 10 kHz using a HEKA EPC 10/2 amplifier with PatchMaster (HEKA Elektronik, Germany). Electrophysiology data were analyzed using PatchMaster and Igor Pro (WaveMetrics, Lake Oswego, Oregon). All mutant fish were genotyped after recording.

### Exploratory screening of ω-conotoxins in ChR-YFP fish

These experiments were performed in the transgenic line expressing ChR-YFP fusion protein primarily in 2°Mn through the a motor neuron-specific GAL4 driver (Gal4^s1020t^; UAS:ChR-YFP, kindly provided by Dr. Claire Wyart, University Pierre et Marie Curie, Paris). Whole field LED illumination was used to excite the 2°Mn using a 475 nm LED driver (Thorlabs, Newton, New Jersey). Whole cell currents coupled to 5 ms pulse of optical excitation given at 10 s intervals were monitored continuously in ventral fast skeletal muscle voltage clamped at −50 mV. Bath solution containing ω-conotoxins (Alomone Labs, Israel) were applied directly to the preparation at indicated concentrations which were based their effective concentrations for the mammalian N-type channels.

### Measurement of dose dependent response for ω–conotoxins on expressed calcium channels

Heterologous expression and recording of zebrafish P/Q and N-type calcium channels was performed as described (Wen et al., 2013). Human embryonic kidney (HEK293T) cells were transfected by Lipofectamine 2000 (Invitrogen, Calsbad, California) with cDNAs for the zebrafish calcium channel α subunit (accession number <u>KC192783</u> for the P/Q channel, <u>KC192784</u> for the N-type), rat α2δ1 subunit (Addgene, accession number <u>AF286488</u>), and the zebrafish β4b subunit (accession number <u>KC192785</u>), in equal molar ratio. To help identifying transfected cells with fluorescence, we used either an EGFP tagged α subunit cDNA or included an empty EGFP vector in the transfection. Cells were re-plated at low density at 24 hrs after transfection and recorded within 48 hrs. Cells with green fluorescence were recorded in the whole-cell voltage-clamp mode with an EPC10/2 amplifier. Patch electrodes (resistances of 3–5 MΩ) contained (in mM): 115 Cs methanesulfonate, 15 CsCl, 5 K-BAPTA, 4 Mg-ATP, and 10 K-HEPES, pH 7.2 with CsOH, ~280 mOsm. The recording bath solution contained (in mM): 134 NaCl, 2.9 KCl, 1.2 MgCl_2_, 2.1 CaCl_2_ and 10 Na-HEPES, pH 7.2, ~290 mOsm. The P/Q channel expressed at reduced levels compared to the N type, requiring substitution of Ba^2+^ for Ca^2+^ as the charge carrier (bath solution containing 10 mM BaCl2, 0.2 mM CaCl_2_). We performed control recordings with Ca^2+^ for comparison. No differences in concentration dependent block were observed between the two charge carriers (n=5). Whole cell currents in response to voltage steps were sampled at 100 kHz and filtered at 10 kHz with PatchMaster acquisition software and leak-subtracted on-line with a p/10 protocol.

For the dose-dependent response, 4 or 5 different concentrations of toxins were applied by a quartz micro-manifold positioned adjacent to the cell (ALA Scientific Instruments, Farmingdale, NY). Since these toxin block are reversible, we alternated with perfusion of bath solution, allowing for full recovery to pre-application level before testing the next concentration. After given sufficient time to reach a steady state block the current-voltage relation for each toxin concentration was determined. Peak currents measured with application of toxins were compared to those with bath solution application.

### Motility analysis

Movement was elicited by a 20 ms 50V stimulus with a Grass SD9 simulator. A stimulus triggered LED light in the field marked the timing of the stimulus. The evoked motility response was recorded at 1000 frames/s using a Fastcam 512-PCI camera (Photron Instruments, San Diego, California). Time course of changes in body curvature was tracked and quantitated using Flote zebrafish motion analysis software (Burgess and Granato, 2007).

### Electron microscopic analysis of the zebrafish NMJ

96-120 hpf wild type zebrafish larvae were high pressure frozen and freeze substituted as described (Helmprobst et al., 2015). Electron micrographs of 250 nm serial sections were recorded at 200 kV at a JEOL JEM-2100 transmission electron microscope with a TVIPS F416 camera (resolution 2048×2048).

### Data analysis and statistical tests

Data were analyzed using PatchMaster and Igor Pro. Statistical analysis was performed with Prism (Graphpad, San Diego, California). Comparison between two samples were done with two-tailed t-test with unequal variances. All data were presented as mean +/− standard deviation (S.D.).

## Acknowledgements

The authors thank Dr. Claudia Lopez from OHSU Multoscale Micrscope Core for her help on electron microscopy, and James Kelly for fish care. We are grateful for support with sample preparation to Marlene Strobel, Daniela Bunsen and Claudia Gehrig-Höhn at the Imaging Core Facility of the Biocenter, University of Würzburg. This work was supported by a grant from NIH to P.B.

